# How convincing is a matching Y-chromosome profile?

**DOI:** 10.1101/131920

**Authors:** Mikkel M Andersen, David J Balding

**Affiliations:** Department of Mathematical Sciences, Aalborg University, Aalborg, Denmark; Section of Forensic Genetics, Department of Forensic Medicine, University of Copenhagen, Copenhagen, Denmark; Centre for Systems Genomics, Royal Parade, University of Melbourne, Vic 3010, Australia; Genetics Institute, University College London, Gower St, WC1E 6BT, UK

## Abstract

The introduction of forensic autosomal DNA profiles was controversial, but the problems were successfully addressed, and DNA profiling has gone on to revolutionise forensic science. Y-chromosome profiles are valuable when there is a mixture of male-source and female-source DNA, and interest centres on the identity of the male source(s) of the DNA. The problem of evaluating evidential weight is even more challenging for Y profiles than for autosomal profiles. Numerous approaches have been proposed, but they fail to deal adequately with the fact that men with matching Y-profiles are related in extended patrilineal clans, many of which may not be represented in available databases. This problem has been exacerbated by recent profiling kits with high mutation rates. Because the relevant population is difficult to define, yet the number of matching relatives is fixed as population size varies, it is typically infeasible to derive population-based match probabilities relevant to a specific crime. We propose a conceptually simple solution, based on a simulation model and software to approximate the distribution of the number of males with a matching Y profile. We show that this distribution is robust to different values for the variance in reproductive success and the population growth rate. We also use importance sampling reweighting to derive the distribution of the number of matching males conditional on a database frequency, finding that this conditioning typically has only a modest impact. We illustrate the use of our approach to quantify the value of Y profile evidence for a court in a way that is both scientifically valid and easily comprehensible by a judge or juror.

## 1 Introduction

A forensic Y-chromosome profile typically consists of the allele at between 15 and 30 short tandem repeat (STR) loci. For autosomal STR profiles, there are two alleles per locus and because of the effects of recombination, the alleles at distinct loci are treated as independent, after any adjustments for sample size, coancestry and direct relatedness. Y profiles usually have only one allele per locus, the exceptions are due to genomic duplications.^1^ The loci lie in the non-recombining part of the Y chromosome, which behaves like a single locus and so the Y profile can be regarded as a single allele, or haplotype. This represents a major contrast with autosomal profiles, the implications of which have not been fully appreciated.^2^

At the core of the weight of evidence for autosomal STR profiles is the match probability, which is the conditional probability that a particular individual *X* has a matching profile, given that the queried contributor Q has it.^3^ Matching at a single, autosomal STR allele is relatively common: typically a few percent of individuals from the same population share a given allele. The probability of matching is increased when X is a relative of Q, but for typical population sizes most of the individuals sharing a given allele are not closely related to Q.

Because of the absence of recombination, the situation for Y-STR profiles is different.^4^ The profile mutation rate is approximately the sum of the mutation rates at each STR locus, which can be above 0.1 per generation for profiling kits in current use (Fig. 1). This, together with the large space of possible profiles, implies that the probability of a matching Y profile between distantly-related males is negligible.^5^ Our results confirm that in practice matches only occur between males connected by a lineage path of at most a few tens of meioses. Thus, Q can have many matching relatives, but the relatedness will typically be too distant to be recognised. Thus an attempt to exclude known patrilineal relatives of Q as alternative sources of the DNA will typically be of limited value. Similarly, while matching surnames can be a guide to matching Y profiles in many societies,^6^ the correlation may not be strong enough for this to be useful in eliminating alternative sources.

**Figure 1.**
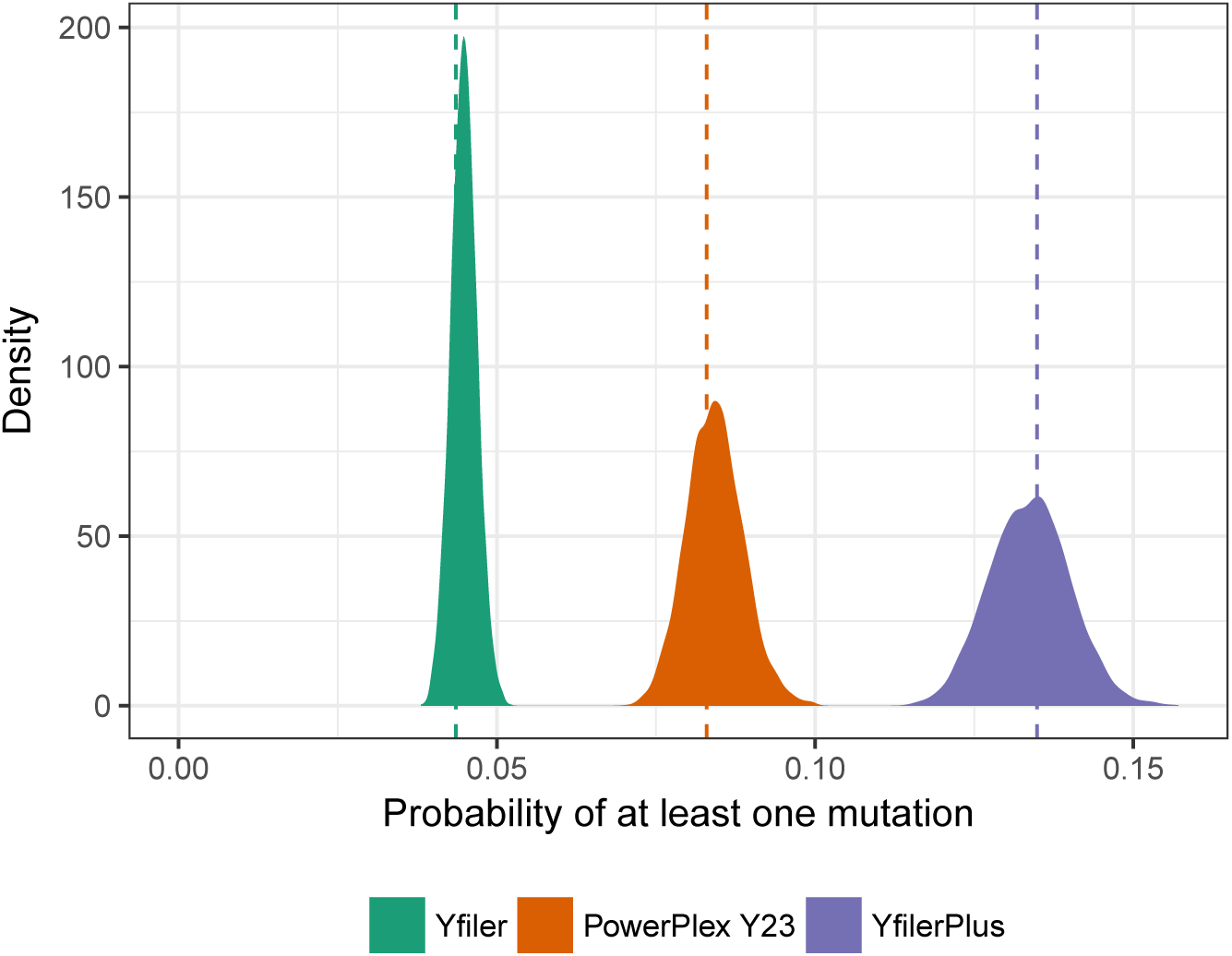
Posterior distributions of the mutation rates. The posterior distributions of the whole-profile per-generation mutation rates for each of three profiling kits: Yfiler, PowerPlex Y23, and Yfiler Plus (see Materials and Methods. Vertical dashed lines indicate maximum-likelihood estimates. See S1 Table for details of the underlying mutation count data.

The number of matching patrilineal relatives of Q is insensitive to population size, and the latter is difficult to define in forensic contexts. Thus, unlike for autosomal profiles, it is difficult to develop an approach to weight-of-evidence based on population match probabilities. Moreover, in most approaches to weight-of-evidence for Y profiles, a count of the profile in a database plays a central role,^7^–^11^ but because of (i) the large number of distinct Y profiles, (ii) the concentration of matching profiles in extended patrilineal clans that may be clustered geographically or socially,^12^ and (iii) the fact that the databases are not scientific random samples,^13^ database information is of limited value, and difficult to interpret in a valid and useful way.^2, 5^

For these reasons we believe that current approaches to evaluating Y profile evidence are unsatisfactory. Proposals have been made that use population genetic models to allow for coancestry,^11, 14^ but the problem remains of setting the coancestry parameter. The appropriate value depends on the fraction of patrilineal relatives of Q in the relevant population, which is not well-defined in most crime cases. Coalescent-based modelling of the match probability can make full use of the available information,^15^ but is computationally demanding and requires the database to be a random sample from the relevant population.

Instead of a population fraction or match probability, we focus on the number of males with Y profile matching that of Q, and we propose a simulation model, implemented in easy-to-use software, to approximate its distribution (Fig. 2). Key parameters of the model include the locus mutation rates, the variance in reproductive success (VRS) and the population growth rate. We extend our simulation framework to provide a novel approach to using counts from a database assumed to be sampled randomly in the relevant population. While this assumption may be unrealistic, it serves to illustrate the limited value of database counts even in this optimistic setting. In contrast with other methods, our approach easily and correctly takes into account a zero database count for the profile of Q.

**Figure 2:**
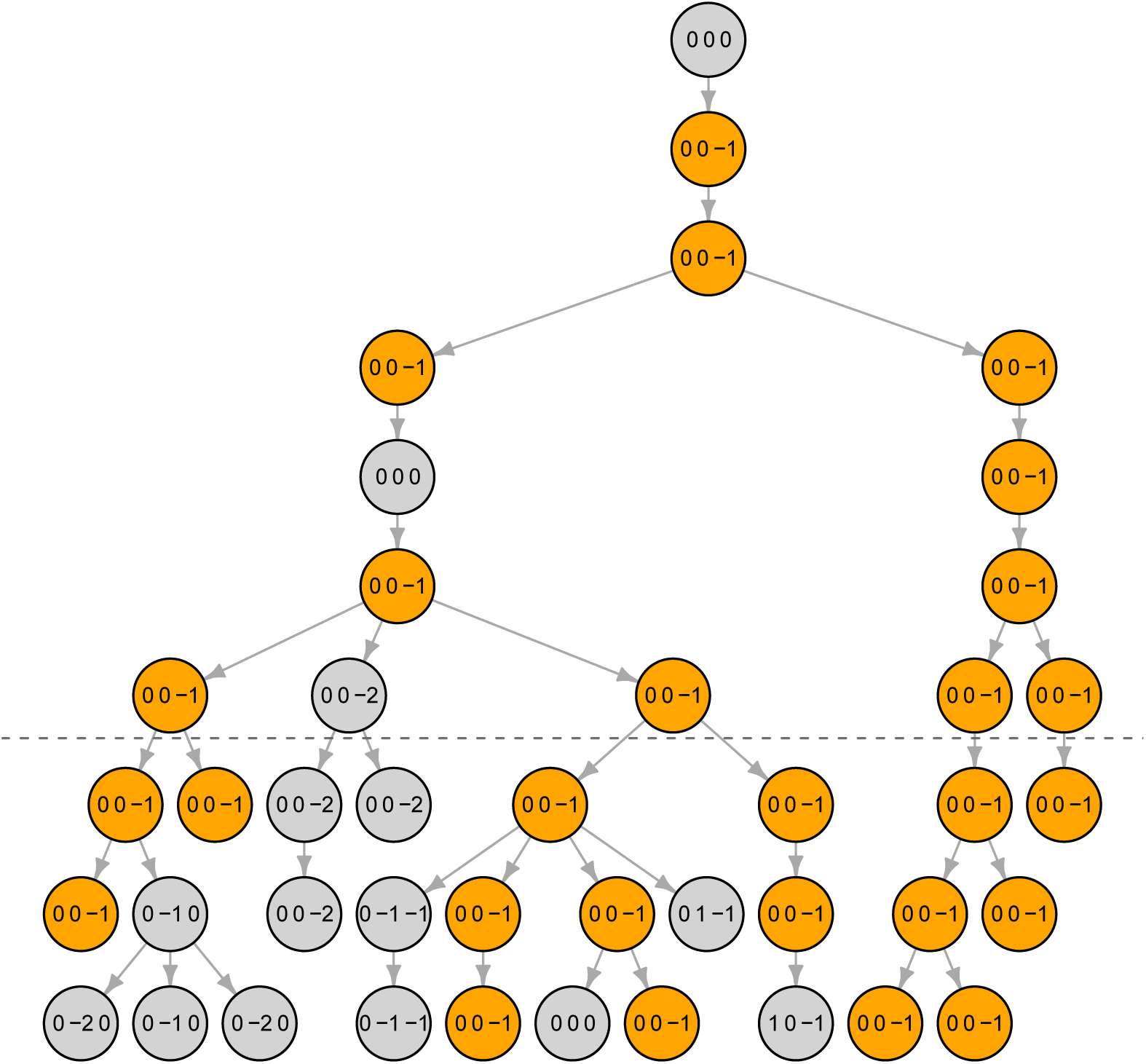
Simulation process illustrated. Simplified illustration of the process of counting haplotype matches (here, 3-locus haplotypes simulated over 10 generations with mutation rate 0.1 per locus per generation). Below the dashed line circles indicate males in the final three generations, who we label “live”. A live male Q was sampled and observed to have haplotype (0, 0*, -*1). The figure shows the founder that is ancestral to Q and all descendants of that founder, highlighting in orange those with haplotype (0, 0*, -*1). In this instance there are 16 live males with haplo-type (0, 0*,-*1); any of these 16 could be Q – the resulting figure would be the same.

Our software and method allow Y profile evidence to be quantified in a way that is valid and directly interpretable to courts.

## 2 Materials and methods

### 2.1 Marker sets and mutation

We consider three Y-STR profiling kits: Yfiler (17 loci), PowerPlex Y23 (23 loci), and Yfiler Plus (27 loci). Mutation count data for these loci are given in S1 Table. All Yfiler loci are present in the other two kits, but PowerPlex Y23 has two loci not present in Yfiler Plus (DYS549 and DYS643) and Yfiler Plus has six loci not present in PowerPlex Y23 (DYS627, DYS460, DYS518, DYS449 and two loci at DYF387S1).

All three kits include both copies of the duplicated locus DYS385. In our simulation study, DYS385a and DYS385b are treated as two independent loci, whereas in practice the order of the duplicated locus is unknown and so e.g. allele pair 13/14 cannot be distinguished from 14/13. We use the same mutation rate for DYS385a and DYS385b, estimated by halving the reported mutation count for locus DYS385. Yfiler Plus includes another duplicated locus, DYF387S1, for which similar comments apply.

### 2.2 Population and mutation simulations

We adopt a Wright-Fisher model, both with a constant per-generation population size *N* and with a growth rate of 2% per generation. Because the profile mutation rate is high, *|*Ω*|*, the number of live males with Y profile matching that of Q, is usually at most several tens, and provided that *N >* 10^3^, it is insensitive to *N*. Although larger than necessary, we chose *N* = 10^5^ for the constant population size, and we verified empirically that increasing this to *N* = 10^6^ led to the same results. The growing population had initial size 7,365 growing over the simulation to 10^6^ in the final generation.

We generate Y profiles for the live individuals by assigning allele 0 at all loci for each founder, and running a neutral, symmetric, single-step mutation process at each locus, forward in time over *G* = 250 generations, so that with high probability all lines of descent from founder to current generation include multiple mutations. We estimate locus mutation rates using the data from S1 Table, allowing for uncertainty by assuming a Beta(1.5, 200) prior distribution (S1 Fig), so that the posterior distribution given the count data is Beta(*x* + 1.5*, y* + 200), where *x* and *y* are the numbers of meioses in which mutations did and did not occur for that locus.

The symmetric, single-step mutation model is a reasonable first approximation to the STR mutation process^16, 17^ and adequate here because most of the matches of interest are between individuals sharing the same haplotype with no intervening mutations, in which case only the mutation rate is important, not any other details of the mutation process. However, some matches arise with intervening mutations that cancel, and so any deficiencies in this model can have a small impact on our results.

The results presented below are based on 5 × 10^5^ values of *|*Ω*|* for each profiling kit, value of VRS and growth rate: we repeated the genealogy simulation 5 times, and for each of these we simulated the mutation process 100 times, each time starting by resampling each mutation rate from its posterior distribution. For each population/mutation simulation we made 10^3^ selections of Q, each time recording the number of his Y-profile matches.

### 2.3 Variable reproductive success

Competition for mates and other factors can lead to a high variance in reproductive success among males in many non-human species and some human societies, notably those in which polygyny is practised.^18^ We perform simulations to investigate the extent to which variability in reproductive success affects the number of Y profiles matching that of Q.

We posit a symmetric Dirichlet distribution that specifies the probability for each man in a generation to be the father of an arbitrary male in the next generation. To avoid dependence on population size, we work with relative paternity probabilities that have mean one and variance denoted VRS. The standard Wright-Fisher model assigns all *N* men a relative probability of paternity equal to one, and VRS = 0. We also consider values 5 and 1 for the parameter of the symmetric Dirichlet, which correspond to VRS = 0.2 and VRS = 1, respectively. When VRS = 0.2, a man’s relative paternity probability has 95% probability to lie between 0.32 and 2.05, and in a constant-size population the corresponding standard deviation in offspring count is about 1.1, which is close to an estimate of 1.07 for the modern US population.^19^ VRS = 1 implies that each man’s relative paternity probability has an Exponential(1) distribution, with 95% equal-tailed interval from 0.025 to 3.7. The corresponding standard deviation in offspring count is about 1.4, which is very high for a modern developed society but higher values have been recorded in human societies.^18^

### 2.4 Comparison of simulated and real databases

The available Y-profile databases are not random samples from a well-defined population. Although sampling biases can also affect databases of autosomal profiles, the mixing effect of recombination reduces their impact, whereas it can be important for Y profiles. We are unable to imic database selection processes, which are diverse, and the populations from which real databases are sampled are often only loosely defined. Therefore we cannot rigorously test the realism of our simulations against empirical data, but some comparison is informative.

We compared results from our population and mutation simulations against data from 6 databases of PowerPlex Y23 profiles drawn from: Central Europe (*n* = 5,361), Eastern Europe (*n* = 1,665), Northern Europe (*n* = 903), Southeastern Europe (*n* = 758), Southern Europe (*n* = 1,462) and Western Europe (*n* = 2,590).^20^ For each real database, we randomly sampled databases with the same *n* from simulated populations with constant population sizes *N* = 10^5^ and *N* = 10^6^. For each profiling kit and value of *N*, we fixed VRS = 0.2 and repeated the population simulation 10 times, for each population we simulated the mutation process 10 times, and for each of these we simulated 10 databases. Hence, each boxplot in Fig. 3 is based on 1,000 datapoints.

**Figure 3:**
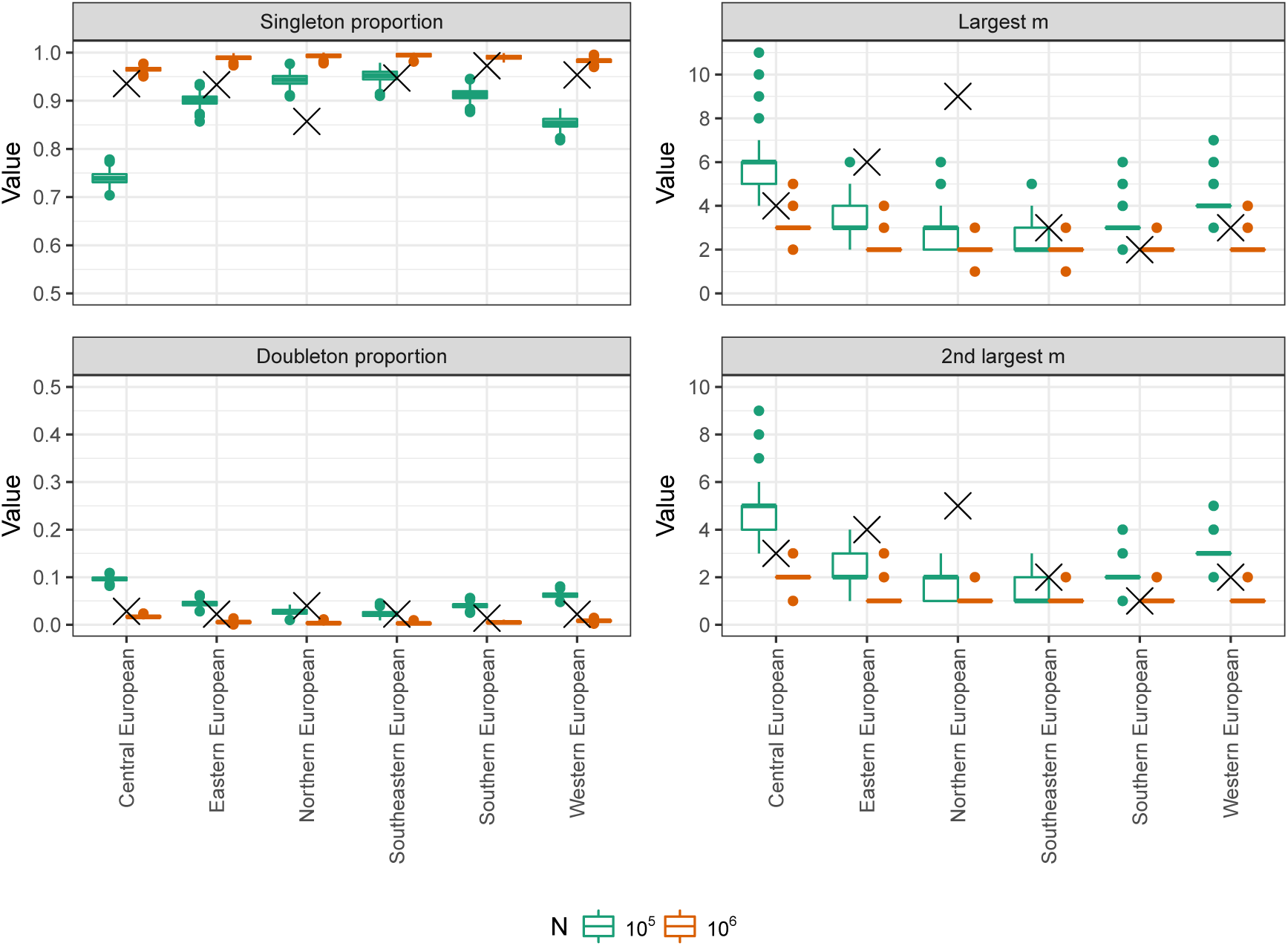
Properties of random databases from simulated populations. Properties of PowerPlex Y23 profile databases drawn at random from simulated populations with VRS = 0.2 and constant population sizes *N* = 10^5^ (green boxplots) and *N* = 10^6^ (red boxplots), and real databases of the same sizes (crosses).^20^ The left panels show the fractions of singleton profiles and doubletons (profiles arising exactly twice). The right panels show the counts of the two most common profiles.

### 2.5 Identifying matching males

Given a population and mutation simulation, we treat the males in the final three generations as potentially of interest, for brevity we label them as “live”. We choose one at random among the live males to represent Q, the queried source of the Y profile and then identify the set Ω of live males carrying the same Y profile as Q. The males in Ω represent the set of potential sources of the crime scene DNA profile, including Q himself. Depending on the case circumstances, some males with Y profiles matching Q will be of an age or live in a location that makes them unlikely to be a source of the crime scene DNA. A forensic DNA expert is not usually entitled to make these judgments, which are a matter for the court and we do not attempt to model such information here.

### 2.6 Importance weighting to condition on database match count

A low database count of the profile suggests a low population count. However because in a large population almost all Y profiles are very rare, a profile count from a database of modest size (typically in practice up to a few thousand) provides little information.^5^ To quantify this, we modify the distribution of *|*Ω*|* by conditioning on a count of *m* copies of the profile of Q in a database of size *n*. Here, we assume that the database has been sampled randomly from the live males, which is typically unrealistic but can be useful for illustration.

We could have obtained the required approximation in our simulation framework as follows: for every simulation of Ω, we sample a database of size *n* at random from the live males, and if the number of males from Ω included in the database equals *m*, we retain the simulation, otherwise we reject it. However, this rejection sampling approach can be inefficient, particularly when *m >* 1.

Instead we use importance sampling to reweight the *|*Ω*|* values to reflect conditioning on *m*. Writing *π* for the fraction of the live individuals that match the profile of Q, the required weights are the binomial probabilities of observing *m* copies of the profile in the database of size *n*:

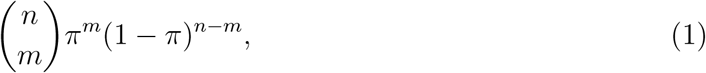

normalised to have an average of one over the full simulation.^21^ Intuitively, a large value of *m* is implausible if *|*Ω*|* is small, and hence *π* is close to zero, resulting in a low weight.

To assess the efficiency of the importance sampling, the effective sample size (ESS) was calculated^21^ for *n* = 100 and 1,000 and for *m* from 0 to 6. The results from the simulations with constant population size (S2 Fig), show that the ESS decreases rapidly as *m* increases, so that the approximation to the conditional distribution of *|*Ω*|* is typically poor for *m >* 3.

## 3 Results

### 3.1 Comparison of simulated and real databases

Fig. 3 shows a broad similarity between real and simulated databases, with properties of real databases often lying between the *N* = 10^5^ and *N* = 10^6^ simulated databases. However these values of *N* are much lower than the census male populations corresponding to the database labels, suggesting restricted sampling. Northern Europe is an outlier, which could reflect non-random sampling. Finland is greatly over-represented in the Northern Europe database relative to its population size, whereas Norway and the Baltic states are not represented at all. Moreover, samples are obtained from just a few centres per country, without comprehensive coverage of the population. In the north of Finland lives one of the most important population isolates in Europe and its over-representation could substantially affect database frequencies. The two most common profiles were both restricted to Finnish samples.

### 3.2 The number of matching Y profiles

Fig. 4 shows, for each of 18 combinations of profiling kit, VRS and constant/growing population size, the distribution of *|*Ω*|*, the number of live males with profile matching that of Q. Here “live” means in the final three generations of the population simulation. As expected, *|*Ω*|* tends to be larger when the haplotype mutation rate is lower. However, the 18 distributions are all highly dispersed and overlap substantially. If Q is a defendant who denies being a source of the DNA, larger values of *|*Ω*|* are more helpful to his case. Therefore, consistent with the recommendation^11^ to use an upper 95% confidence interval for the population frequency, if a court wishes to consider scenarios that are helpful to the defendant while remaining realistic, it might regard the 95% quantile as a useful summary of the distribution of *|*Ω*|* (Table 1).

**Figure 4:**
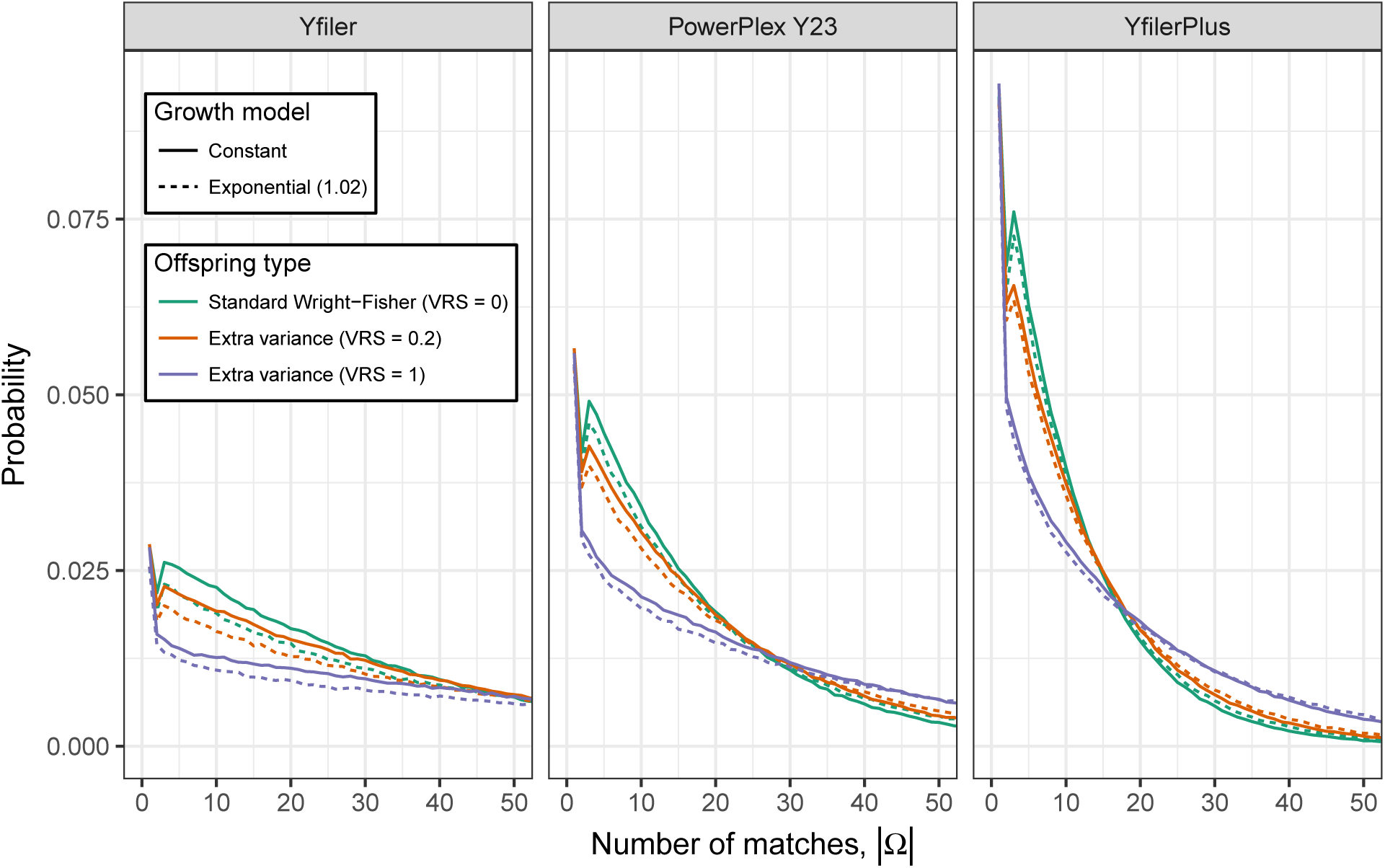
The distribution of *|*Ω*|*, the number of live males with Y profile matching that of Q. The distribution is shown for each of three profiling kits, three values of the variance in reproductive success (VRS) and with and without population growth (growth rates 1 and 1.02). See Table 1 for numerical properties of these distributions.

**Table 1:**
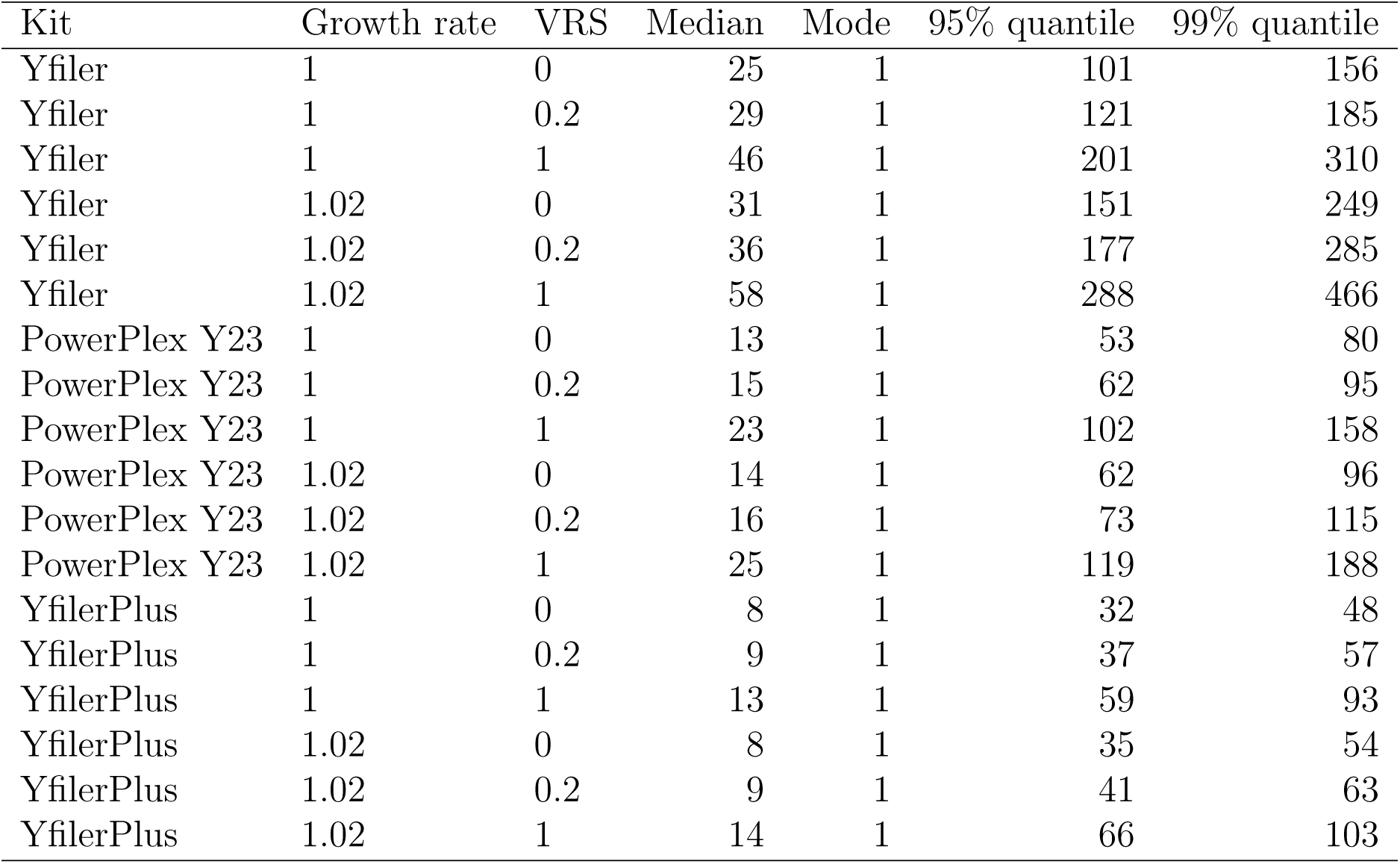
**Properties of the unconditional distribution of** *|*Ω*|* **the number of males with a Y profile matching that of Q.** Each row corresponds to a distinct combination of profiling kit, value of VRS and growth model.

Fig. 5 shows for the same 18 parameter combinations as Fig. 4, the distribution of Δ, the number of meioses between Q and another male with matching Y profile. We see that matching males are predominantly separated from Q by a handful of meioses (such as uncles and cousins), but there exist matches separated by *>* 20 meioses. While this is too remote for the relatedness to be recognised, almost all matching males are more closely related to Q than is typical for pairs drawn from the population, which implies that “random man” match probabilities are misleading for Y profiles.

**Figure 5:**
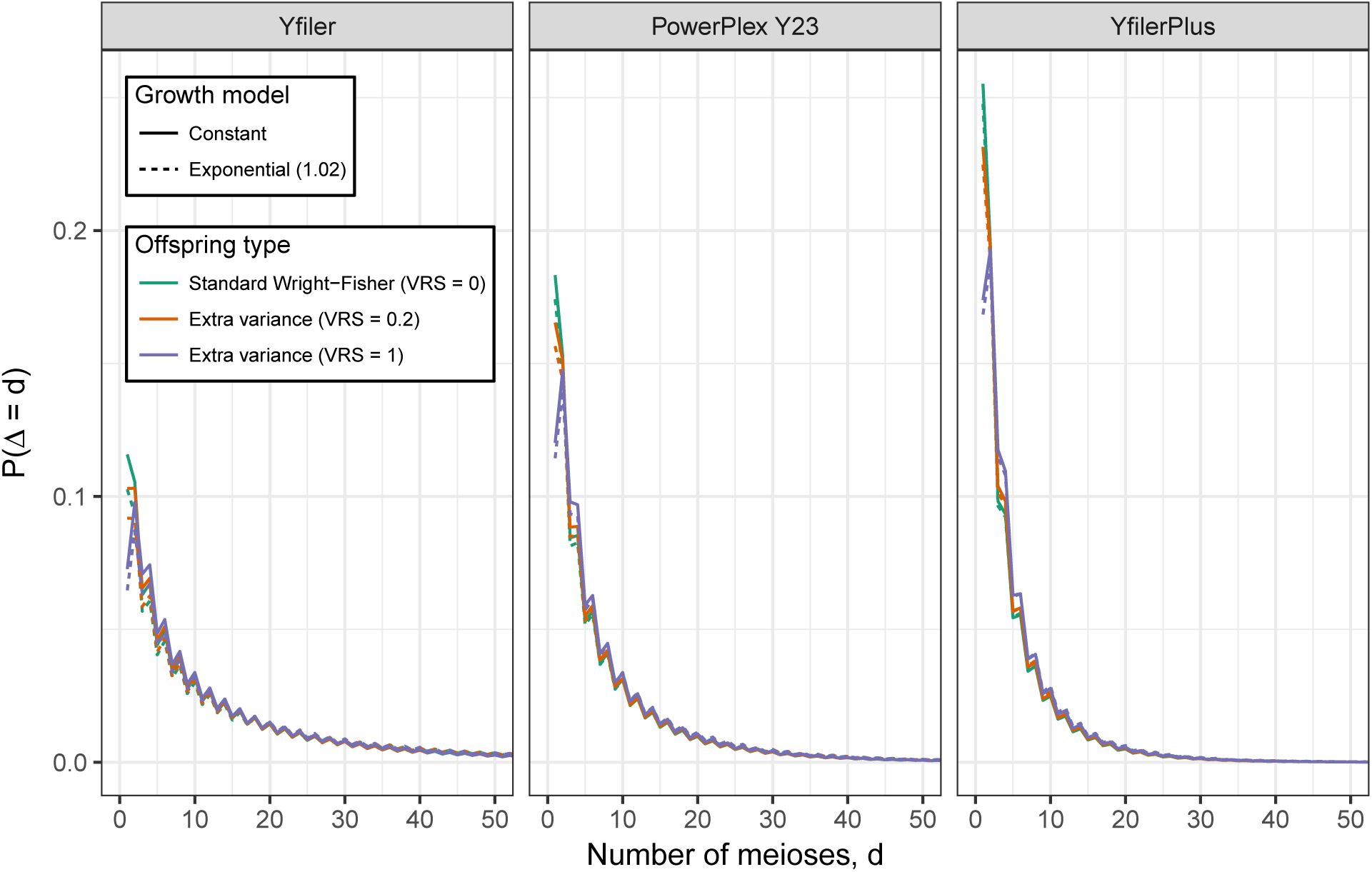
The distribution of Δ, the number of father-son steps between Q and another male in Ω. The distribution is shown for each of three profiling kits, three values of VRS and with and without population growth. See S2 Table for numerical properties of these distributions.

From Fig. 5 we can infer that Y profile matches between distantly-related males are so unlikely as to be negligible. In the limit as their common ancestor is increasingly further back in the past (that is, as Δ increases), the match probability converges to the product of population allele fractions, which is negligibly small for a full Y profile from a modern kit. Our simulations exaggerate this possibility because all founders 250 generations in the past are assigned the same haplotype. Nevertheless for our most discriminating kit, YfilerPlus, no matches were observed between any pair of descendants of distinct founders over a total of 4.5 million population simulations.

### 3.3 Role of database information

Table 2 gives properties of the conditional distribution of *|*Ω*|* given three values of the database count *m*, for each of three database sizes *n*, drawn from a simulated constant-size population with VRS = 0.2. As expected, *|*Ω*|* increases stochastically with *m*, but the distributions substantially overlap and the practical impact of the dependence on *m* for the decision process of a court may be only modest. For example, with the most discriminatory kit that we consider, Yfiler Plus, the upper 95% quantile of the unconditional distribution of Ω is 37 (Table 2, row 1). If, however, we note that the profile of Q is unobserved in a database of size *n* = 10^3^, this apparently useful information only has the effect of reducing the 95% quantile from 37 to 36, even under optimistic assumptions about the database. In reality, the males in *|*Ω*|* are likely to be clustered geographically and/or socially, and the database is usually not a random sample from the population, further reducing the value of database information.

**Table 2:** Properties of the distribution of the number of males with a matching Yfiler Plus profile with and without database count information. Row 1 is the same as the fifth last row of Table 1 (YfilerPlus, constant population size, VRS = 0.2; “-” indicates the absence of *m* and *n* values). Other rows show properties of the corresponding conditional distributions given an observation of *m* copies of the profile in a randomly-sampled database of size *n*. Corresponding values for other profiling kits, VRS values, and population growth models are given in S3 Table through S8 Table.

| m | n | Median | Mode | 95% quantile | 99% quantile |
| --- | --- | --- | --- | --- | --- |
| - | - | 9 | 1 | 37 | 57 |
| 0 | 100 | 9 | 1 | 37 | 57 |
| 0 | 1,000 | 9 | 1 | 36 | 55 |
| 0 | 10,000 | 6 | 1 | 27 | 41 |
| 1 | 100 | 21 | 11 | 58 | 81 |
| 1 | 1,000 | 20 | 11 | 56 | 78 |
| 1 | 10,000 | 15 | 9 | 41 | 58 |
| 2 | 100 | 33 | 27 | 77 | 102 |
| 2 | 1,000 | 32 | 21 | 74 | 99 |
| 2 | 10,000 | 23 | 17 | 55 | 73 |

Properties of conditional distributions for other VRS values and growth models are given in S3 Table (VRS = 0.2, constant population size), S4 Table (VRS = 1, constant population size), S5 Table (VRS = 0, constant population size), S6 Table (VRS = 0.2, population growth), S7 Table (VRS = 1, population growth), S8 Table (VRS = 0, population growth).

## 4 Discussion

### 4.1 Presentation in court

Although the distribution of *|*Ω*|* varies with demographic parameters such as VRS and population growth rate, which cannot be known exactly for a specific court case, we have shown (Fig. 4) that the distributions are insensitive to major changes in parameter values, relative to the precision necessary for a juror’s reasoning: whether the number of matching individuals in the population is 40 or 50 or 60 is unlikely to have much impact on a juror’s decision, but orders of magnitude may well be important.

Based on the results from simulations such as those underlying Fig. 4, an expert could propose a suitable summary of the distribution of *|*Ω*|* over a range of plausible parameter values, and this could be presented at court along the following lines:

> “A Y-chromosome profile was recovered from the crime scene. Mr Q has a matching Y profile and so is not excluded as a contributor of DNA. Using population genetics theory and data, we conclude that the number of males in the population with a matching Y profile is probably less than 20, and is very unlikely (probability *<* 5%) to exceed 40. These men or boys span a wide range of ages and we don’t know where they live. They are all paternal-line relatives of Q, but the relationship may extend over many father-son steps, well beyond the known relatives of Q. Since these individuals share paternal-line ancestry with Q, they could also be similar to Q in ethnic identity, language, religion, physical appearance and place of residence.”

If there is database frequency information, the court could be further advised, for example:

> “The Y profile of Q was not observed in a database of 1,000 profiles. Because the database does not represent a scientific random sample and because paternal-line relatives may tend to be clustered in geographic and social groups that are not well sampled in the database, it is difficult to interpret this information. If the database were a random sample from the population, its effect would be to reduce the 95% upper limit on the number of matching males from 40 to 39.”

The impact of this information may be minimal, and it could perhaps be omitted except that courts may be expecting to hear database information. Depending on the circumstances of the case, a judge might further instruct members of the jury:

> “If you consider that there may be up to 40 males of different ages with a Y profile matching that of Q, and that these males may tend to resemble Q in some characteristics more than random members of the population, your task is to decide whether all the evidence that has been presented to you is enough to convince you that Q is the source of the crime scene DNA, and not one of these other males with the same Y profile.”

It should be clear from these considerations that a matching Y profile, taken alone, can never suffice to establish convincingly that Q is a source of the crime-scene DNA. However, it remains very powerful evidence that can reduce the number of alternative sources from perhaps several millions, for a crime in a large city, down to just a few tens. If an autosomal DNA profile is also available that includes Q then the remaining task for the prosecution of establishing that Q is the source, rather than one of the 40 or so potential Y-matchers, will usually be readily achievable even if the autosomal profile is complex, for example due to DNA from multiple sources. Alternatively, non-DNA evidence can suffice to complete the task of convincing a jury that Q is the source of the Y profile.

We have here only considered DNA samples with a single male contributor. A similar approach can be applied for multiple male contributors, see **??**, S3 Fig and S9 Table.

### 4.2 Coancestry, kinship and sampling adjustments

For autosomal DNA profiles a coancestry adjustment is generally recommended to allow for remote shared ancestry between Q and X, the hypothetical alternative source of the DNA. Similarly if Q and X might be close relatives, a match probability can be computed using explicit kinship parameters. Sampling adjustments are also sometimes recommended, based on database size. None of these adjustments is needed for the method proposed here, because all matches with related individuals are modelled in the simulations, as is the size of the database.

### 4.3 Likelihood ratio reporting

Forensic weight-of-evidence is often best quantified using a ratio of likelihoods under prosecution and defence hypotheses, which in simple settings reduces to a match probability. We support that approach in general, but there are specific difficulties applying it to Y profiles, because the match probability for an alternative to Q will depend strongly on Δ, the number of meioses separating that individual from Q. We therefore propose a different approach, reporting to the court an estimate of the number of males with matching Y profiles. Estimating the number of matching individuals in the population has been recommended in the past for autosomal DNA profiles. In the mid-1990s, when autosomal DNA profile match probabilities were not as minuscule as they have become, the England and Wales Court of Appeal recommended that, instead of a match probability, courts be informed of the expected number of matching individuals in a relevant population^3^ (see Section 11.4.3). This recommendation was followed for some time, until autosomal match probabilities became too small for the approach to be helpful to jurors. In addition, for complex profiles the concept of the number of matching individuals is problematic, because the “match” may only be partial.

For older Y-profiling kits with lower profile mutation rate, or when only a few loci generate usable results due to poor sample quality, it may be appropriate to use a standard match probability approach, because in that case there will be many matching individuals in the population, so that profile population fractions are larger and so can be better estimated from databases, with sampling biases being less important than when matching individuals are predominantly closely related to Q.

### 4.4 Mitochondrial DNA profiles

Despite their lack of discriminatory power relative to autosomal or even Y profiles, mitochondrial DNA (mtDNA) profiles can be invaluable when matrilineal relatedness is of interest or when nuclear DNA is unavailable, for example in old bones, teeth or hair shafts, or for some highly-degraded samples.^22^ Similar issues arise for mtDNA profiles as for Y profiles, but because the mutation rate for the entire mtDNA genome is an order of magnitude lower than for current Y profiling kits, the sets of matching individuals tend to be much larger. This makes the problems of relatedness of matching individuals and database sampling bisases less severe, though still important.

### 4.5 Software

Our open source (Apache License) R^23^ package malan (MAle Lineage ANlysis) available at https://github.com/mikldk/malan performed the analyses described in this paper. A vignette demonstrating the functionality of malan is available in the package. We also provide an online demonstration app based on malan but with limited functionality, available at <u>https://mikldk.shinyapps.io/ychr-matches/</u>. Mutation count data are provided for the Yfiler, PowerPlex 23 and Yfiler Plus kits.

## 4.6 Acknowledgments

The authors wish to thank the Isaac Newton Institute for Mathematical Sciences, Cambridge UK, for support and hospitality during the programme Probability and Statistics in Forensic Science, where this paper was conceived. The programme was supported by EPSRC grant no EP/K032208/1. We thank Francois Balloux (UCL), Poul Svante Eriksen (AAU) and Niels Morling (UCPH) for helpful discussions.

## A Supplementary Text: Mixtures

Crime-scene Y profiles can reflect a mixture of DNA from multiple male sources. It may be possible to identify the profiles underlying the mixture based on differing amounts of DNA or similarity to observed profiles,^9^, ^24^ but often they cannot be fully deconvolved with high confidence. We also performed simulations of a mixed Y profile with DNA from two male sources. For each of the three values of VRS and the Yfiler Plus profiling kit, we performed 10 population simulations, for each of these 10 mutation simulations were performed, and for each of these 1000 pairs of males were chosen at random to be the contributors to the mixture. For each contributor pair, the number of included live individuals was recorded, where “included” means has a Y profile consisting only of alleles observed in the mixture.

Among the included individuals will be those with profiles exactly matching one of the reference individuals. The sizes of these two sets of individuals will be approximately independent and have the same distribution as *|*Ω*|* for a single-source Y profile described above. S3 Fig. and S9 Table show that the number of included males who do not exactly match either reference individual is stochastically smaller than *|*Ω*|*. Therefore the distribution of the total number of included males is conservatively estimated as the sum of three independent versions of *|*Ω*|*. Further work is required to develop procedures for presenting evidence in court based on mixed Y profiles that takes available database information into account, but our simulations will provide helpful guidance.

## B Supplementary figures

**S1 Fig.**
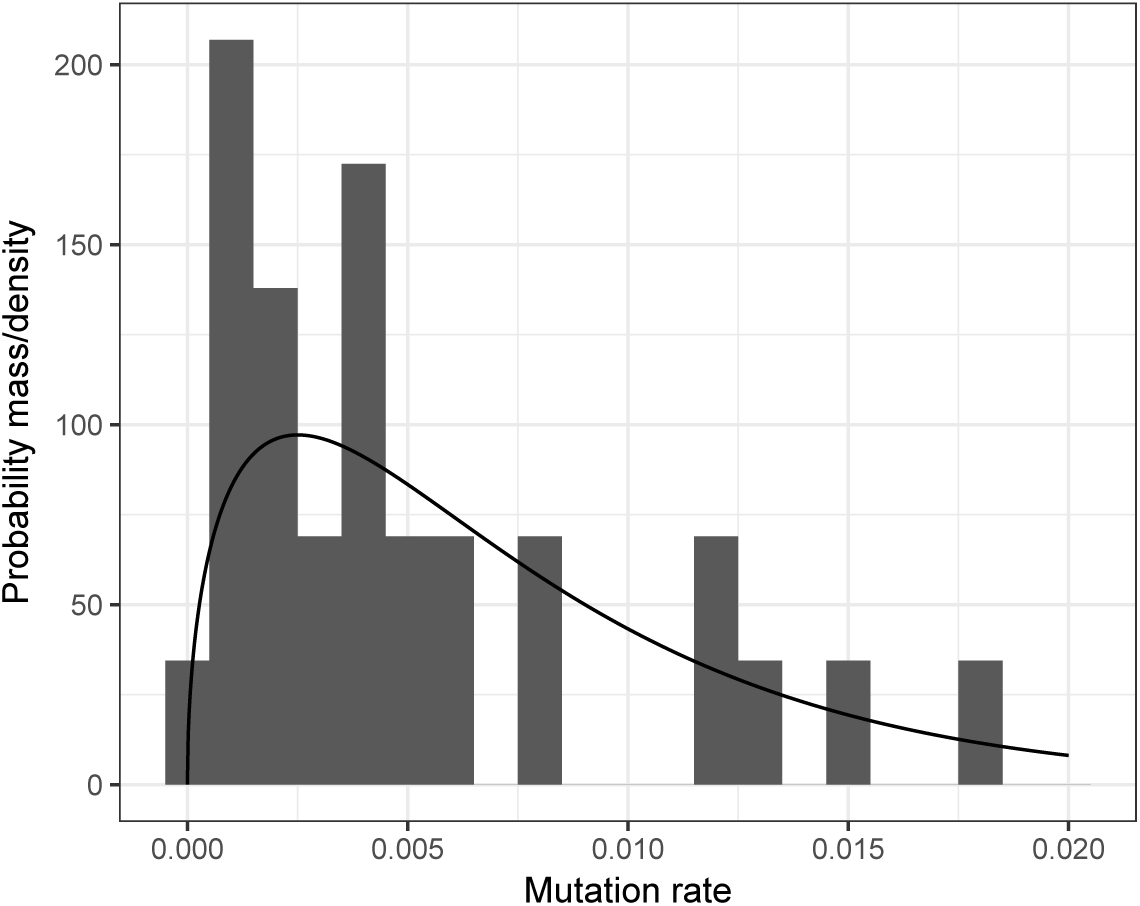
Y-STR mutation rates. Histogram bars show empirical mutation rates for the 29 loci included in the three Y-STR profiling kits (rates obtained as a ratio of the counts in S1HYPERLINK \l "bookmark36" Table). The two duplicated loci (DYS385 and DYF387S1) are each represented as two loci with the same mutation rate. The curve shows the probability density for the Beta(1.5, 200) prior distribution assumed for each mutation rate.

**S2 Fig.**
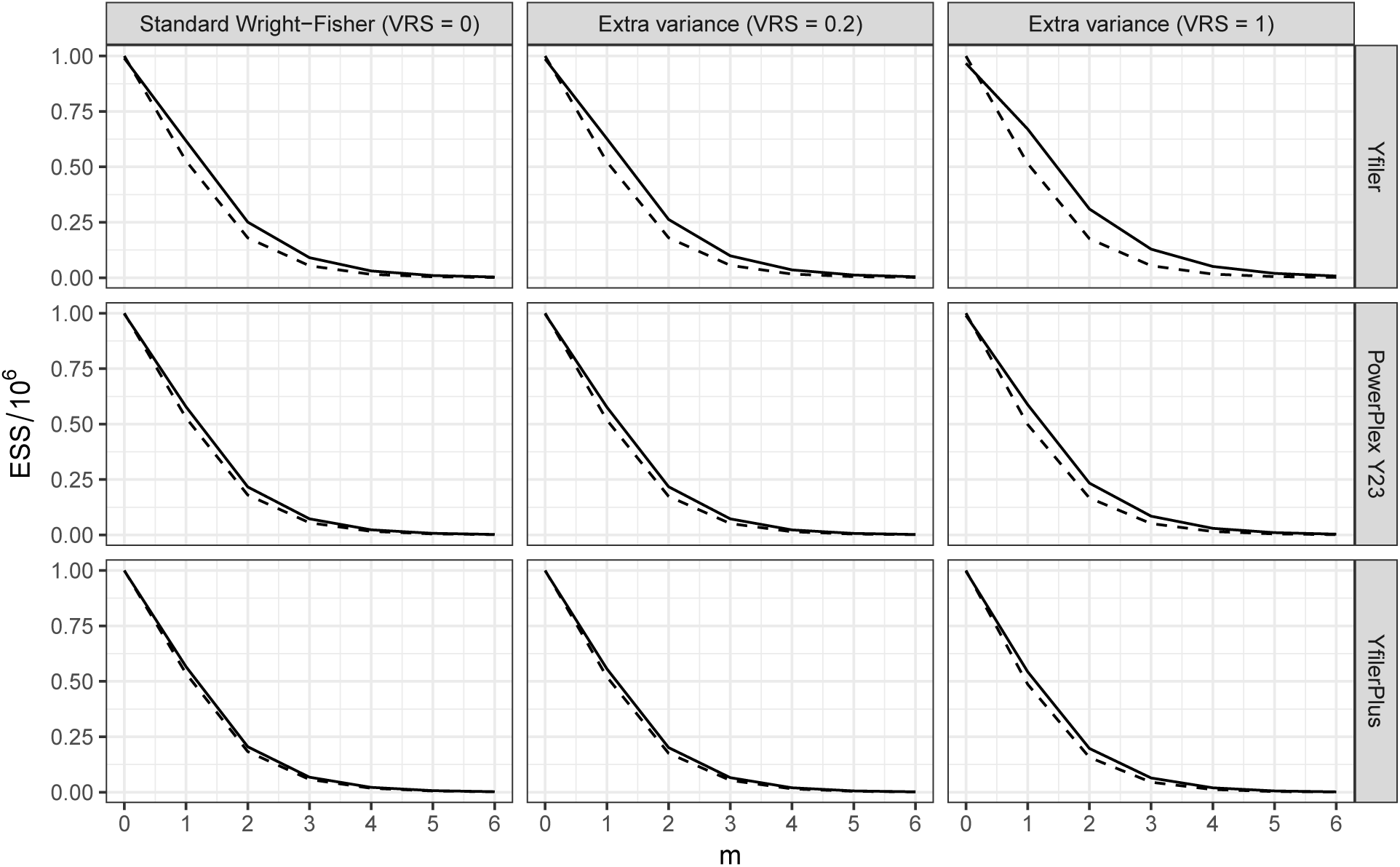
Importance sampling ESS. The effective sample size for simulations with constant population size, as a fraction of the 10^6^ simulated *|*Ω*|* values, for the importance sampling to approximate distributions for *|*Ω*|* conditional on database profile count *m*, for *m* from 0 to 6. The database sizes are *n* = 100 (dashed line) and *n* = 1,000 (solid line).

**S3 Fig.**
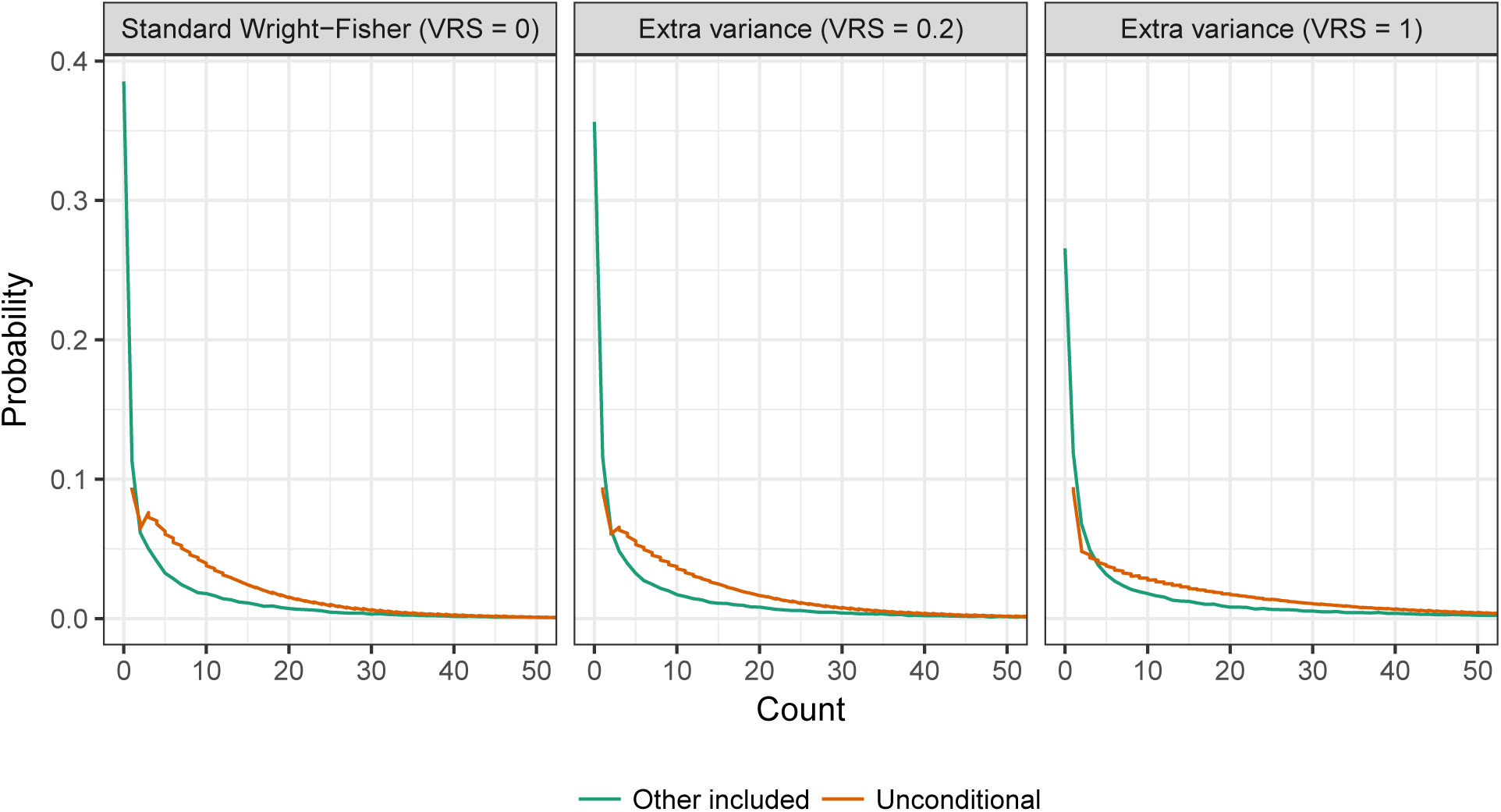
Count distribution for a two-person mixture. Distribution of the number of Y profiles included in a mixed Yfiler Plus profile arising from two male contributors. The red curve corresponds to profiles that exactly match the profile of one of the contributors. It is the same as the red (“unconditional”) curve in Fig. 4 and is included again here for comparison. The green curve corresponds to “other included”: males with a Y profile that consist entirely of alleles within the profiles of the two contributors, and which therefore cannot be excluded from being one of the contributing profiles, but they do not fully match either of the two contributing profiles.

## C Supplementary tables

**S1 Table.**
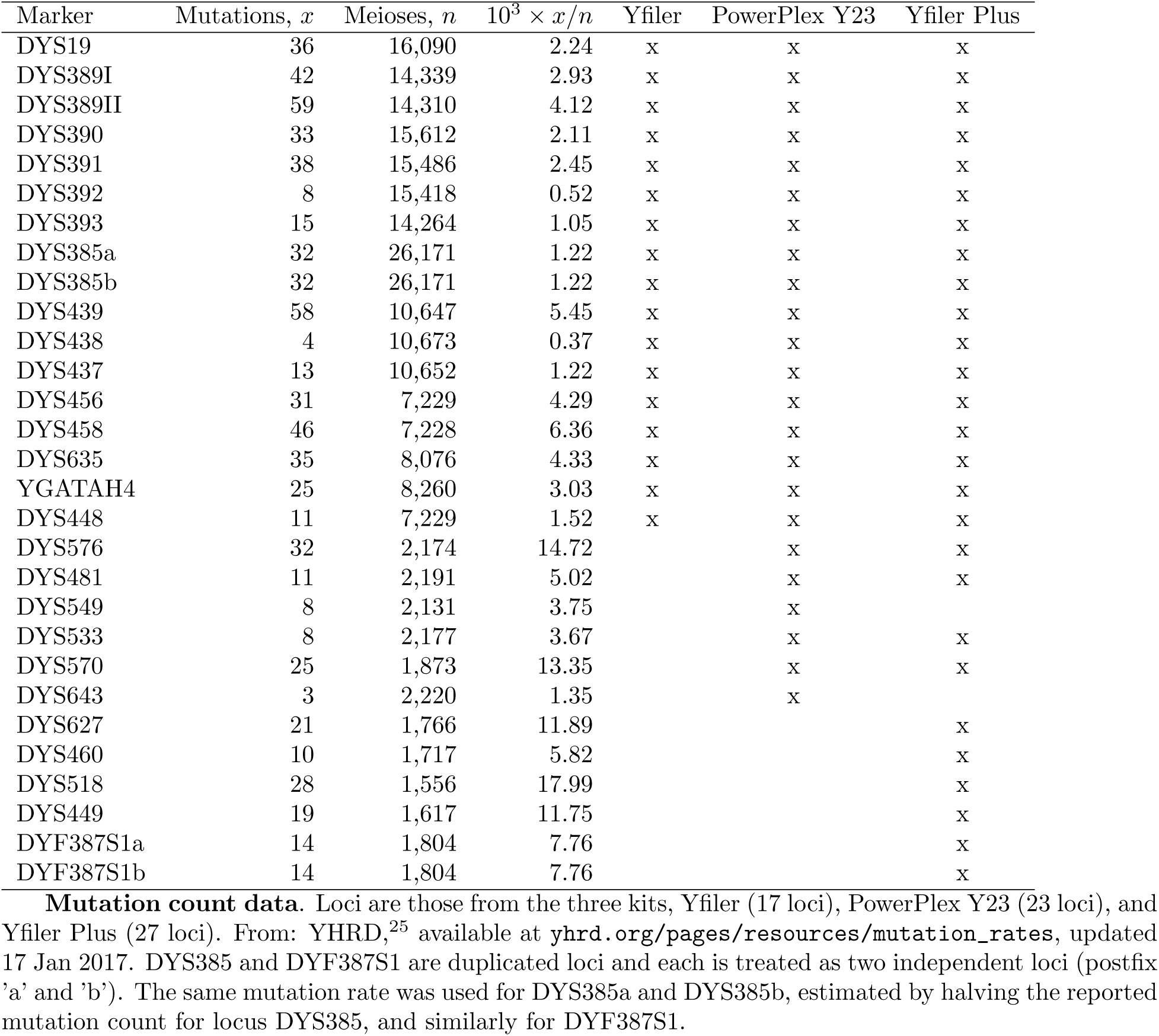
Mutation count data. Loci are those from the three kits, Yfiler (17 loci), PowerPlex Y23 (23 loci), and Yfiler Plus (27 loci). From: YHRD,^25^ available at yhrd.org/pages/resources/mutation_rates, updated 17 Jan 2017. DYS385 and DYF387S1 are duplicated loci and each is treated as two independent loci (postfix ‘a’ and ‘b’). The same mutation rate was used for DYS385a and DYS385b, estimated by halving the reported mutation count for locus DYS385, and similarly for DYF387S1.

| Marker | Mutations, $x$ | Meioses, $n$ | $10^3 \times x/n$ | Yfiler | PowerPlex Y23 | Yfiler Plus |
| --- | --- | --- | --- | --- | --- | --- |
| DYS19 | 36 | 16,090 | 2.24 | x | x | x |
| DYS389I | 42 | 14,339 | 2.93 | x | x | x |
| DYS389II | 59 | 14,310 | 4.12 | x | x | x |
| DYS390 | 33 | 15,612 | 2.11 | x | x | x |
| DYS391 | 38 | 15,486 | 2.45 | x | x | x |
| DYS392 | 8 | 15,418 | 0.52 | x | x | x |
| DYS393 | 15 | 14,264 | 1.05 | x | x | x |
| DYS385a | 32 | 26,171 | 1.22 | x | x | x |
| DYS385b | 32 | 26,171 | 1.22 | x | x | x |
| DYS439 | 58 | 10,647 | 5.45 | x | x | x |
| DYS438 | 4 | 10,673 | 0.37 | x | x | x |
| DYS437 | 13 | 10,652 | 1.22 | x | x | x |
| DYS456 | 31 | 7,229 | 4.29 | x | x | x |
| DYS458 | 46 | 7,228 | 6.36 | x | x | x |
| DYS635 | 35 | 8,076 | 4.33 | x | x | x |
| YGATAH4 | 25 | 8,260 | 3.03 | x | x | x |
| DYS448 | 11 | 7,229 | 1.52 | x | x | x |
| DYS576 | 32 | 2,174 | 14.72 |  | x | x |
| DYS481 | 11 | 2,191 | 5.02 |  | x | x |
| DYS549 | 8 | 2,131 | 3.75 |  | x |  |
| DYS533 | 8 | 2,177 | 3.67 |  | x | x |
| DYS570 | 25 | 1,873 | 13.35 |  | x | x |
| DYS643 | 3 | 2,220 | 1.35 |  | x |  |
| DYS627 | 21 | 1,766 | 11.89 |  |  | x |
| DYS460 | 10 | 1,717 | 5.82 |  |  | x |
| DYS518 | 28 | 1,556 | 17.99 |  |  | x |
| DYS449 | 19 | 1,617 | 11.75 |  |  | x |
| DYF387S1a | 14 | 1,804 | 7.76 |  |  | x |
| DYF387S1b | 14 | 1,804 | 7.76 |  |  | x |

**S2 Table.**
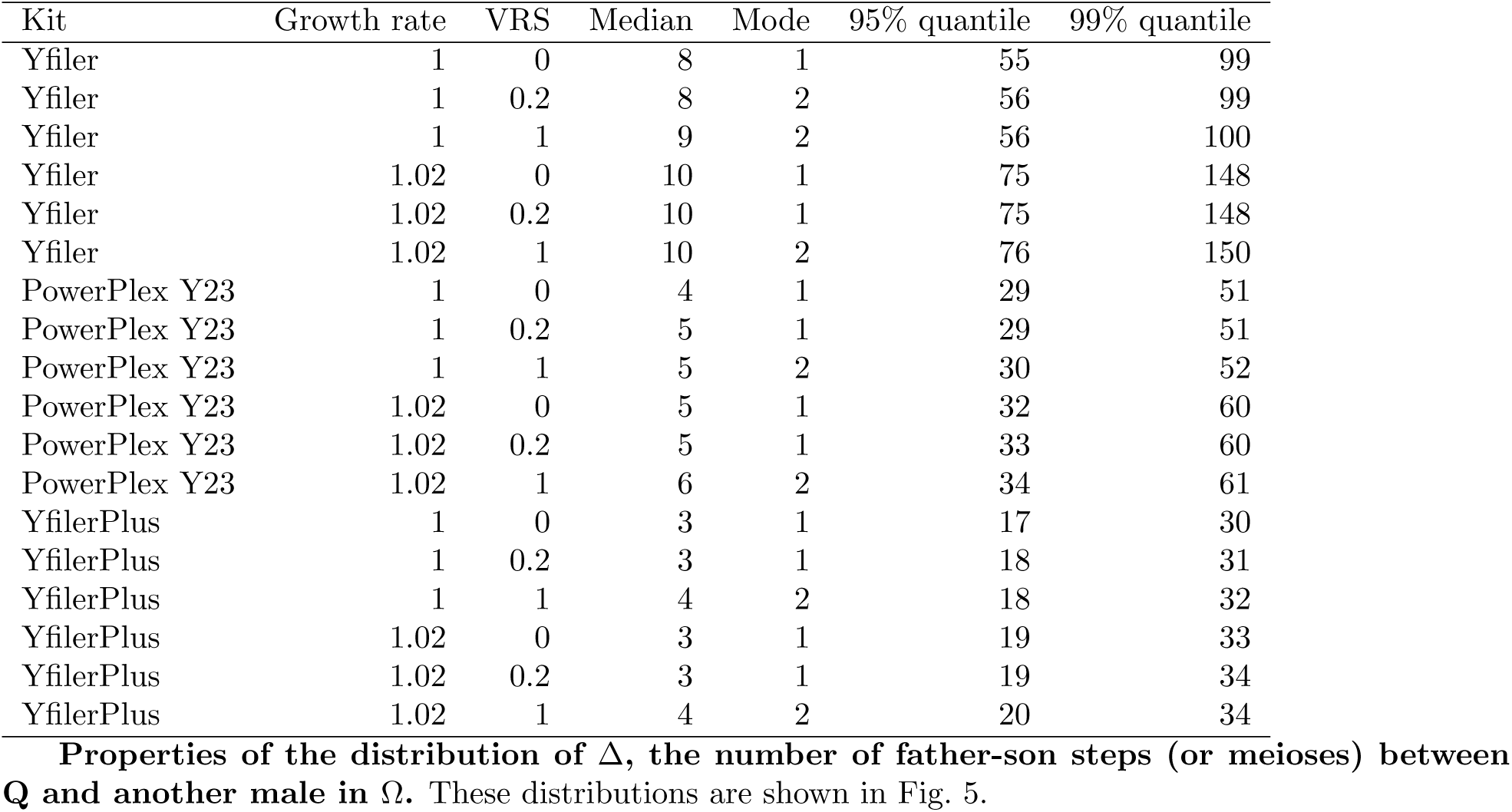
**Properties of the distribution of** Δ**, the number of father-son steps (or meioses) between Q and another male in** Ω. These distributions are shown in Fig. 5.

**S3 Table.** The number of matching live individuals for VRS = 0.2 and constant population size. Highlighted in grey are key properties of the unconditional distributions of *|*Ω*|*, the number of live individuals with profile matching that of Q, shown in Fig. 4. Other rows show corresponding properties of conditional distributions given an observation of *m* copies of the profile in a randomly-sampled database of size *n*. “-” means unconditional in the sense that database profile counts (*m*) are not taken into account. The YfilerPlus results are shown in Table 2, and repeated here for comparison with the other two profiling kits.

| Kit | m | n | Median | Mode | 95% quantile | 99% quantile |
| --- | --- | --- | --- | --- | --- | --- |
| Yfiler | - | - | 29 | 1 | 121 | 185 |
| Yfiler | 0 | 100 | 28 | 1 | 119 | 183 |
| Yfiler | 0 | 1,000 | 25 | 1 | 107 | 163 |
| Yfiler | 0 | 10,000 | 13 | 1 | 52 | 80 |
| Yfiler | 1 | 100 | 67 | 41 | 188 | 261 |
| Yfiler | 1 | 1,000 | 60 | 35 | 168 | 235 |
| Yfiler | 1 | 10,000 | 29 | 16 | 82 | 114 |
| Yfiler | 2 | 100 | 106 | 72 | 247 | 329 |
| Yfiler | 2 | 1,000 | 95 | 72 | 222 | 295 |
| Yfiler | 2 | 10,000 | 46 | 35 | 108 | 144 |
| PowerPlex Y23 | - | - | 15 | 1 | 62 | 95 |
| PowerPlex Y23 | 0 | 100 | 15 | 1 | 61 | 94 |
| PowerPlex Y23 | 0 | 1,000 | 14 | 1 | 58 | 89 |
| PowerPlex Y23 | 0 | 10,000 | 9 | 1 | 37 | 57 |
| PowerPlex Y23 | 1 | 100 | 34 | 19 | 97 | 136 |
| PowerPlex Y23 | 1 | 1,000 | 32 | 19 | 91 | 128 |
| PowerPlex Y23 | 1 | 10,000 | 21 | 13 | 58 | 81 |
| PowerPlex Y23 | 2 | 100 | 54 | 39 | 129 | 174 |
| PowerPlex Y23 | 2 | 1,000 | 51 | 36 | 121 | 163 |
| PowerPlex Y23 | 2 | 10,000 | 33 | 27 | 76 | 102 |
| YfilerPlus | - | - | 9 | 1 | 37 | 57 |
| YfilerPlus | 0 | 100 | 9 | 1 | 37 | 57 |
| YfilerPlus | 0 | 1,000 | 9 | 1 | 36 | 55 |
| YfilerPlus | 0 | 10,000 | 6 | 1 | 27 | 41 |
| YfilerPlus | 1 | 100 | 21 | 11 | 58 | 81 |
| YfilerPlus | 1 | 1,000 | 20 | 11 | 56 | 78 |
| YfilerPlus | 1 | 10,000 | 15 | 9 | 41 | 58 |
| YfilerPlus | 2 | 100 | 33 | 27 | 77 | 102 |
| YfilerPlus | 2 | 1,000 | 32 | 21 | 74 | 99 |
| YfilerPlus | 2 | 10,000 | 23 | 17 | 55 | 73 |

**S4 Table.** Similar to S3 Table but with VRS = 1. Please see S3 Table caption for details.

| Kit | m | n | Median | Mode | 95% quantile | 99% quantile |
| --- | --- | --- | --- | --- | --- | --- |
| Yfiler | - | - | 46 | 1 | 201 | 310 |
| Yfiler | 0 | 100 | 45 | 1 | 197 | 303 |
| Yfiler | 0 | 1,000 | 37 | 1 | 164 | 253 |
| Yfiler | 0 | 10,000 | 14 | 1 | 61 | 95 |
| Yfiler | 1 | 100 | 110 | 68 | 313 | 442 |
| Yfiler | 1 | 1,000 | 92 | 45 | 261 | 367 |
| Yfiler | 1 | 10,000 | 35 | 21 | 98 | 137 |
| Yfiler | 2 | 100 | 177 | 129 | 422 | 572 |
| Yfiler | 2 | 1,000 | 147 | 116 | 348 | 467 |
| Yfiler | 2 | 10,000 | 55 | 45 | 130 | 174 |
| PowerPlex Y23 | - | - | 23 | 1 | 102 | 158 |
| PowerPlex Y23 | 0 | 100 | 22 | 1 | 100 | 156 |
| PowerPlex Y23 | 0 | 1,000 | 20 | 1 | 91 | 141 |
| PowerPlex Y23 | 0 | 10,000 | 10 | 1 | 47 | 72 |
| PowerPlex Y23 | 1 | 100 | 56 | 39 | 162 | 226 |
| PowerPlex Y23 | 1 | 1,000 | 51 | 29 | 147 | 205 |
| PowerPlex Y23 | 1 | 10,000 | 26 | 16 | 75 | 106 |
| PowerPlex Y23 | 2 | 100 | 91 | 65 | 215 | 288 |
| PowerPlex Y23 | 2 | 1,000 | 82 | 65 | 196 | 259 |
| PowerPlex Y23 | 2 | 10,000 | 42 | 29 | 100 | 135 |
| YfilerPlus | - | - | 13 | 1 | 59 | 93 |
| YfilerPlus | 0 | 100 | 13 | 1 | 59 | 92 |
| YfilerPlus | 0 | 1,000 | 12 | 1 | 56 | 87 |
| YfilerPlus | 0 | 10,000 | 8 | 1 | 35 | 55 |
| YfilerPlus | 1 | 100 | 33 | 20 | 97 | 137 |
| YfilerPlus | 1 | 1,000 | 31 | 20 | 91 | 129 |
| YfilerPlus | 1 | 10,000 | 20 | 11 | 57 | 80 |
| YfilerPlus | 2 | 100 | 54 | 39 | 131 | 172 |
| YfilerPlus | 2 | 1,000 | 51 | 37 | 123 | 163 |
| YfilerPlus | 2 | 10,000 | 32 | 26 | 76 | 103 |

**S5 Table.** Similar to S3 Table but with VRS = 0. The population size is constant so this case corresponds to a standard Wright-Fisher model. Please see S3 Table caption for details.

| Kit | m | n | Median | Mode | 95% quantile | 99% quantile |
| --- | --- | --- | --- | --- | --- | --- |
| Yfiler | - | - | 25 | 1 | 101 | 156 |
| Yfiler | 0 | 100 | 24 | 1 | 100 | 154 |
| Yfiler | 0 | 1,000 | 22 | 1 | 91 | 140 |
| Yfiler | 0 | 10,000 | 12 | 1 | 49 | 74 |
| Yfiler | 1 | 100 | 56 | 34 | 158 | 221 |
| Yfiler | 1 | 1,000 | 51 | 30 | 143 | 201 |
| Yfiler | 1 | 10,000 | 27 | 15 | 75 | 105 |
| Yfiler | 2 | 100 | 88 | 66 | 210 | 283 |
| Yfiler | 2 | 1,000 | 81 | 66 | 190 | 254 |
| Yfiler | 2 | 10,000 | 42 | 30 | 99 | 133 |
| PowerPlex Y23 | - | - | 13 | 1 | 53 | 80 |
| PowerPlex Y23 | 0 | 100 | 13 | 1 | 52 | 80 |
| PowerPlex Y23 | 0 | 1,000 | 12 | 1 | 50 | 76 |
| PowerPlex Y23 | 0 | 10,000 | 8 | 1 | 34 | 51 |
| PowerPlex Y23 | 1 | 100 | 29 | 18 | 81 | 113 |
| PowerPlex Y23 | 1 | 1,000 | 27 | 16 | 77 | 107 |
| PowerPlex Y23 | 1 | 10,000 | 19 | 10 | 52 | 73 |
| PowerPlex Y23 | 2 | 100 | 46 | 33 | 107 | 144 |
| PowerPlex Y23 | 2 | 1,000 | 44 | 33 | 102 | 136 |
| PowerPlex Y23 | 2 | 10,000 | 29 | 21 | 69 | 92 |
| YfilerPlus | - | - | 8 | 1 | 32 | 48 |
| YfilerPlus | 0 | 100 | 8 | 1 | 32 | 48 |
| YfilerPlus | 0 | 1,000 | 8 | 1 | 31 | 47 |
| YfilerPlus | 0 | 10,000 | 6 | 1 | 24 | 36 |
| YfilerPlus | 1 | 100 | 17 | 10 | 49 | 67 |
| YfilerPlus | 1 | 1,000 | 17 | 10 | 47 | 66 |
| YfilerPlus | 1 | 10,000 | 13 | 9 | 37 | 51 |
| YfilerPlus | 2 | 100 | 27 | 22 | 64 | 85 |
| YfilerPlus | 2 | 1,000 | 27 | 20 | 62 | 83 |
| YfilerPlus | 2 | 10,000 | 21 | 16 | 48 | 64 |

**S6 Table.** Similar to S3 Table but with VRS = 0.2 and population size growth. The population growth rate is 2% per generation. Please see S3 Table caption for details.

| Kit | m | n | Median | Mode | 95% quantile | 99% quantile |
| --- | --- | --- | --- | --- | --- | --- |
| Yfiler | - | - | 36 | 1 | 177 | 285 |
| Yfiler | 0 | 100 | 36 | 1 | 176 | 284 |
| Yfiler | 0 | 1,000 | 35 | 1 | 173 | 278 |
| Yfiler | 0 | 10,000 | 30 | 1 | 144 | 229 |
| Yfiler | 1 | 100 | 98 | 50 | 309 | 474 |
| Yfiler | 1 | 1,000 | 96 | 50 | 301 | 458 |
| Yfiler | 1 | 10,000 | 80 | 50 | 245 | 356 |
| Yfiler | 2 | 100 | 171 | 128 | 473 | 763 |
| Yfiler | 2 | 1,000 | 167 | 128 | 453 | 724 |
| Yfiler | 2 | 10,000 | 136 | 89 | 344 | 488 |
| PowerPlex Y23 | - | - | 16 | 1 | 73 | 115 |
| PowerPlex Y23 | 0 | 100 | 16 | 1 | 73 | 115 |
| PowerPlex Y23 | 0 | 1,000 | 16 | 1 | 73 | 114 |
| PowerPlex Y23 | 0 | 10,000 | 15 | 1 | 67 | 106 |
| PowerPlex Y23 | 1 | 100 | 41 | 23 | 120 | 171 |
| PowerPlex Y23 | 1 | 1,000 | 40 | 23 | 119 | 169 |
| PowerPlex Y23 | 1 | 10,000 | 37 | 23 | 110 | 156 |
| PowerPlex Y23 | 2 | 100 | 67 | 54 | 162 | 219 |
| PowerPlex Y23 | 2 | 1,000 | 66 | 54 | 161 | 219 |
| PowerPlex Y23 | 2 | 10,000 | 61 | 41 | 149 | 201 |
| YfilerPlus | - | - | 9 | 1 | 41 | 63 |
| YfilerPlus | 0 | 100 | 9 | 1 | 41 | 63 |
| YfilerPlus | 0 | 1,000 | 9 | 1 | 41 | 63 |
| YfilerPlus | 0 | 10,000 | 9 | 1 | 39 | 60 |
| YfilerPlus | 1 | 100 | 23 | 15 | 65 | 91 |
| YfilerPlus | 1 | 1,000 | 23 | 15 | 65 | 90 |
| YfilerPlus | 1 | 10,000 | 22 | 13 | 62 | 87 |
| YfilerPlus | 2 | 100 | 37 | 31 | 86 | 114 |
| YfilerPlus | 2 | 1,000 | 37 | 26 | 86 | 113 |
| YfilerPlus | 2 | 10,000 | 35 | 24 | 83 | 110 |

**S7 Table.** Similar to S3 Table but with VRS = 1 and population size growth. The population growth rate is 2% per generation. Please see S3 Table caption for details.

| Kit | m | n | Median | Mode | 95% quantile | 99% quantile |
| --- | --- | --- | --- | --- | --- | --- |
| Yfiler | - | - | 58 | 1 | 288 | 466 |
| Yfiler | 0 | 100 | 58 | 1 | 287 | 464 |
| Yfiler | 0 | 1,000 | 56 | 1 | 277 | 447 |
| Yfiler | 0 | 10,000 | 43 | 1 | 209 | 334 |
| Yfiler | 1 | 100 | 161 | 86 | 502 | 737 |
| Yfiler | 1 | 1,000 | 155 | 86 | 483 | 707 |
| Yfiler | 1 | 10,000 | 117 | 57 | 356 | 514 |
| Yfiler | 2 | 100 | 276 | 169 | 717 | 1049 |
| Yfiler | 2 | 1,000 | 266 | 169 | 686 | 993 |
| Yfiler | 2 | 10,000 | 198 | 169 | 494 | 675 |
| PowerPlex Y23 | - | - | 25 | 1 | 119 | 188 |
| PowerPlex Y23 | 0 | 100 | 25 | 1 | 119 | 187 |
| PowerPlex Y23 | 0 | 1,000 | 25 | 1 | 117 | 185 |
| PowerPlex Y23 | 0 | 10,000 | 22 | 1 | 104 | 164 |
| PowerPlex Y23 | 1 | 100 | 67 | 43 | 198 | 281 |
| PowerPlex Y23 | 1 | 1,000 | 66 | 43 | 195 | 278 |
| PowerPlex Y23 | 1 | 10,000 | 59 | 29 | 172 | 245 |
| PowerPlex Y23 | 2 | 100 | 110 | 80 | 269 | 359 |
| PowerPlex Y23 | 2 | 1,000 | 109 | 80 | 266 | 355 |
| PowerPlex Y23 | 2 | 10,000 | 96 | 80 | 233 | 314 |
| YfilerPlus | - | - | 14 | 1 | 66 | 103 |
| YfilerPlus | 0 | 100 | 14 | 1 | 66 | 103 |
| YfilerPlus | 0 | 1,000 | 14 | 1 | 65 | 102 |
| YfilerPlus | 0 | 10,000 | 13 | 1 | 61 | 95 |
| YfilerPlus | 1 | 100 | 37 | 22 | 108 | 154 |
| YfilerPlus | 1 | 1,000 | 37 | 22 | 107 | 153 |
| YfilerPlus | 1 | 10,000 | 34 | 22 | 100 | 142 |
| YfilerPlus | 2 | 100 | 60 | 53 | 146 | 195 |
| YfilerPlus | 2 | 1,000 | 60 | 53 | 145 | 194 |
| YfilerPlus | 2 | 10,000 | 56 | 44 | 135 | 182 |

**S8 Table.** Similar to S3 Table but with VRS = 0 and population size growth. The population growth rate is 2% per generation. Please see S3 Table caption for details.

| Kit | m | n | Median | Mode | 95% quantile | 99% quantile |
| --- | --- | --- | --- | --- | --- | --- |
| Yfiler | - | - | 31 | 1 | 151 | 249 |
| Yfiler | 0 | 100 | 31 | 1 | 151 | 248 |
| Yfiler | 0 | 1,000 | 30 | 1 | 148 | 243 |
| Yfiler | 0 | 10,000 | 26 | 1 | 126 | 203 |
| Yfiler | 1 | 100 | 84 | 44 | 274 | 438 |
| Yfiler | 1 | 1,000 | 83 | 44 | 267 | 421 |
| Yfiler | 1 | 10,000 | 70 | 44 | 218 | 324 |
| Yfiler | 2 | 100 | 149 | 119 | 460 | 883 |
| Yfiler | 2 | 1,000 | 146 | 102 | 435 | 774 |
| Yfiler | 2 | 10,000 | 120 | 78 | 318 | 468 |
| PowerPlex Y23 | - | - | 14 | 1 | 62 | 96 |
| PowerPlex Y23 | 0 | 100 | 14 | 1 | 62 | 96 |
| PowerPlex Y23 | 0 | 1,000 | 14 | 1 | 61 | 95 |
| PowerPlex Y23 | 0 | 10,000 | 13 | 1 | 58 | 90 |
| PowerPlex Y23 | 1 | 100 | 34 | 22 | 100 | 142 |
| PowerPlex Y23 | 1 | 1,000 | 34 | 22 | 99 | 141 |
| PowerPlex Y23 | 1 | 10,000 | 32 | 17 | 93 | 132 |
| PowerPlex Y23 | 2 | 100 | 56 | 41 | 135 | 179 |
| PowerPlex Y23 | 2 | 1,000 | 55 | 41 | 134 | 178 |
| PowerPlex Y23 | 2 | 10,000 | 52 | 41 | 125 | 168 |
| YfilerPlus | - | - | 8 | 1 | 35 | 54 |
| YfilerPlus | 0 | 100 | 8 | 1 | 35 | 54 |
| YfilerPlus | 0 | 1,000 | 8 | 1 | 35 | 54 |
| YfilerPlus | 0 | 10,000 | 8 | 1 | 34 | 52 |
| YfilerPlus | 1 | 100 | 19 | 9 | 56 | 77 |
| YfilerPlus | 1 | 1,000 | 19 | 9 | 55 | 77 |
| YfilerPlus | 1 | 10,000 | 18 | 9 | 53 | 75 |
| YfilerPlus | 2 | 100 | 31 | 25 | 74 | 99 |
| YfilerPlus | 2 | 1,000 | 31 | 25 | 74 | 98 |
| YfilerPlus | 2 | 10,000 | 30 | 25 | 71 | 94 |

**S9 Table.** Mixed profiles. Key properties of the distribution of the number of Y profiles included in a mixed Yfiler Plus profile arising from two male contributors in a constant-size population. See S3 Fig legend for explanations of “other included” and “unconditional”.

| VRS | Type | Median | Mode | 95% quantile | 99% quantile |
| --- | --- | --- | --- | --- | --- |
| 0 | Other included | 2 | 0 | 30 | 53 |
| 0 | Unconditional | 8 | 1 | 32 | 48 |
| 0.2 | Other included | 2 | 0 | 35 | 63 |
| 0.2 | Unconditional | 9 | 1 | 37 | 57 |
| 1 | Other included | 3 | 0 | 56 | 102 |
| 1 | Unconditional | 13 | 1 | 59 | 93 |

## References

1 JM Butler, AE Decker, MC Kline, and PM Vallone. Chromosomal duplications along the Y chromosome and their potential impact on Y-STR interpretation. J Forensic Sci, 50(4):853–9, 2005.

2 P de Knijff. Son, give up your gun: Presenting Y-STR results in court. Profiles in DNA, 6(2):3–5, 2003.

3 C.D. Steele and D. Balding. Weight of evidence for forensic DNA proflles. Wiley, 2nd ed. edition, 2015.

4 A Caliebe, A Jochens, S Willuweit, L Roewer, and M Krawczak. No shortcut solution to the problem of Y-STR match probability calculation. Forensic Science International: Genetics, 15:69–75, 2015.

5 C Brenner. Understanding Y haplotype matching probability. Forensic Science International: Genetics, 8:233–43, 2014.

6 T King and M Jobling. What’s in a name? y chromosomes, surnames and the genetic genealogy revolution. Trends in Genetics, 25(8):351–60, 2009.

7 Charles H. Brenner. Fundamental problem of forensic mathematics – The evidential value of a rare haplotype. Forensic Science International: Genetics, 4(5):281–291, 2010.

8 Mikkel Meyer Andersen, Amke Caliebe, Arne Jochens, Sascha Willuweit, and Michael Krawczak. Estimating trace-suspect match probabilities for singleton Y-STR haplotypes using coalescent theory. Forensic Science International: Genetics, 7:264–271, 2013.

9 Mikkel Meyer Andersen, Poul Svante Eriksen, and Niels Morling. The discrete Laplace exponential family and estimation of Y-STR haplotype frequencies. Journal of Theoretical Biology, 329:39–51, 2013.

10 Giulia Cereda. Impact of model choice on LR assessment in case of rare haplotype match (frequentist approach). Scandinavian Journal of Statistics, (to appear), 2017.

11 Scientific Working Group on DNA Analysis Methods. Interpretation Guidelines for YChromosome STR Typing. Available at https://www.swgdam.org/, pages 9–15, 2014.

12 L Roewer. Y chromosome STR typing in crime casework. Forensic Sci Med Pathol, 5:77– 84, 2009.

13 P Gill, C Brenner, B Brinkmann, B Budowle, A Carracedo, and M Jobling. DNA Commission of the International Society of Forensic Genetics: recommendations on forensic analysis using Y-chromosome STRs. Forensic Science International, 124:5–10, 2001.

14 J Buckleton, M Krawczak, and B Weir. The interpretation of lineage markers in forensic DNA testing. Forensic Science International: Genetics, 5(78-83), 2011.

15 I Wilson, M Weale, and D Balding. Inferences from DNA data: population histories, evolutionary processes and forensic match probabilities. J Roy Statist Soc A, 166(2):155– 188, 2003.

16 F Hao and J Chu. A brief review of short tandem repeat mutation. Genomics, Proteomics & Bioinformatics, 5(1):7–14, 2007.

17 K Ballantyne, M Goedbloed, R Fang, O Schaap, O Lao, A Wollstein, Y Choi, K van Duijn, M Vermeulen, S Brauer, M Furtado, and M Kayser. Mutability of Y-chromosomal microsatellites: Rates, characteristics, molecular bases, and forensic implications. The American Journal of Human Genetics, 87:341–353, 2010. 17

18 GR Brown, KN Laland, and MB Mulder. Bateman’s principles and human sex roles. Trends Ecol Evol, 24(6-14):297–304, 2009.

19 J Weeden, MJ Abrams, MC Green, and J Sabini. Do high-status people really have fewer children? Hum Nat, 17:377, 2006.

20 J Purps, S Siegert, S Willuweita, M Nagya, C Alvesc, R Salazar, and S Angustia. A global analysis of Y-chromosomal haplotype diversity for 23 STR loci. Forensic Science International: Genetics, 12:12–23, 2014.

21 Augustine Kong. A Note on Importance Sampling using Standardized Weights. Technical Report 348, Department of Statistics, University of Chicago, 1992. https://galton.uchicago.edu/techreports/tr348.pdf.

22 W Parson, L Gusmaô, D Hares, J Irwin, W Mayr, N Morling, E Pokorak, M Prinz, A Salas, P Schneider, and T Parsons. DNA Commission of the International Society of Forensic Genetics: Revised and extended guidelines for mitochondrial DNA typing. Forensic Science International: Genetics, 13:134–142, 2014.

23 R Core Team. R: A Language and Environment for Statistical Computing. R Foundation for Statistical Computing, Vienna, Austria, 2017.

24 Mikkel Meyer Andersen, Poul Svante Eriksen, Helle Smidt Mogensen, and Niels Morling. Identifying the most likely contributors to a Y-STR mixture using the discrete Laplace method. Forensic Science International: Genetics, 15:76–83, 2015.

25 S Willuweit and L Roewer. The new Y chromosome haplotype reference database. Forensic Science International: Genetics, 15:43–48, 2015.

